# A synthetic communication system uncovers extracellular immunity that self-limits bacteriophage transmission

**DOI:** 10.1101/2022.05.11.491355

**Authors:** Amit Pathania, Corbin Hopper, Amir Pandi, Matthias Függer, Thomas Nowak, Manish Kushwaha

## Abstract

Understanding how delivery and exchange of genetic information by bacteriophages shapes bacterial populations is important for designing applications for phage therapy, biocontrol, and microbiome engineering. Here, we present a synthetic intercellular communication system that repurposes phage M13 for genetic exchange between *Escherichia coli* cells and build mathematical models of the communication behaviour. Our models, based on Chemical Reaction Networks, capture the growth burden, cell density, and growth phase dependence of phage secretion and infection kinetics and predict the stochasticity characterising phage-bacterial interactions at low numbers. In co-cultures of phage sender and receiver cells, resource sharing and selection pressure determine the choice of horizontal versus vertical phage transmission. Surprisingly, we discover that a phage-encoded immunity factor confers extracellular protection to uninfected bacteria, reducing infection rates by 70%. In a simulated gut environment, this novel “self-jamming” mechanism enables the phage to farm uninfected bacteria for future infections, increasing the overall success of both M13 and *E. coli*. The synthetic system developed here lays the groundwork for implementing population level controls in engineered bacterial communities, using phage signals for communication.

## Introduction

Bacteriophages are estimated to be the most abundant biological entities on the planet^1,2^. They play key roles in shaping microbial communities by not only controlling bacterial numbers but also disseminating a wide range of genetically encoded functions among members of those communities^3–5^, including antibiotic resistance^6^, sulphur metabolism^7^, bioremediation^8,9^, virulence^10^, and immune escape^11^. The ability of phages to package and deliver genetic material has been leveraged for several applications ranging from phage therapy against infectious agents^12,13^, curing of antimicrobial resistance genes^14^, and editing of microbiomes^15,16^, to delivery of metabolic pathways^17^, phage display^18^, and continuous evolution systems^19,20^. Phages used for these applications exhibit a diversity of infection cycles with respect to their bacterial hosts, broadly classified as lytic, lysogenic, or chronic^3^, depending on whether the infection cycle is productive and whether it results in cellular lysis.

While lytic phages are generally preferred for phage therapy and biocontrol applications^21^, concerns around endotoxin release during bacterial lysis have prompted a search for alternative non-lytic strategies^22^, including the use of chronic phages. Chronic phages have been used to deliver regulatory RNAs to silence antibiotic resistance genes in bacteria^23^, edit microbiomes^15^, or even kill targeted bacteria by non-lytic means^12,22^. A majority of chronic phages belong to the Inoviridae family that is widespread in a variety of environments including the human gut^24^, where they constitute ∼20% of the phageome^25^. These filamentous phages play key roles in their natural ecosystems^26^, affecting pathogenicity^10,11^, biofilm formation^27^, motility and chemotaxis^28^ of their hosts. Their non-lytic infection cycle and modest growth burden for the host also make them amenable to easier plasmid-like manipulation. Consequently, they have found uses in many applications like nanotechnology^29^, phage display^18^, phage therapy^12^, microbiome editing^15^, vaccine development^30^, biosensing^31^, and directed evolution^19,20^.

Used extensively in many biotechnology applications, M13 and its other Ff relatives are the most commonly studied filamentous phages^32^. Their infection kinetics has been studied^33–35^ and modelled for decades. Available models are deterministic, either tracking intracellular molecular concentrations within an infected cell^36,37^, or phenomenologically describing population-level densities of infected cells over time^38^. The early mathematical models are based on data from natural phage infections, where the packaging and infection functions are encoded on the same DNA molecule; so, secretion and infection are concurrent. However, most applications of M13 use phagemids, where the packaging signal is located on a different DNA molecule than the rest of the infection machinery^39^. As a result, the secretion and infection processes are often spatially and temporally separated. Furthermore, current infection models typically consider the log phase of bacterial growth with constant infection and secretion rates, ignoring the effect of growth phase on the secretion and infection processes^33^. This makes them unrealistic for simulations of M13 behaviour in most complex environments where growth conditions of *E. coli* vary widely^40^. To effectively engineer the M13 machinery for delivery of DNA, comprehensive models of their secretion and infection kinetics are needed.

Here, we repurpose the phage M13 machinery for intercellular exchange of genetic information, building and characterising the synthetic communication system using a series of experiments with increasing complexity (Fig. S1). Sender cells secrete DNA messages as packaged phagemids, carrying a bacterial replication origin and the M13 packaging signal (PS)^41^. They also contain the M13 helper plasmid that encodes the functional protein machinery but no PS. The messages can then infect receiver cells carrying the F-plasmid, which encodes the receptor required for efficient M13 entry^37,42^. We build, parameterise, and integrate realistic quantitative Chemical Reaction Network (CRN) models of the communication system from 52192 experimental data-points. The models capture the growth burden of maintaining the communication machinery, the relationship between cellular growth phase and the sending and receiving kinetics of phage signals, as well as the effect of antibiotic resistance. In contrast to the existing deterministic models, the CRN models allow for both stochastic and deterministic simulations. We then show that the stochasticity included in our CRN models is key to predicting experimental outcomes from low-frequency infection events. Interestingly, the stochasticity of infection can be reduced by using sender cells instead of isolated phages for infection. When using conditioned media to study the effect of nutrient depletion in long-term experiments, we surprisingly found a significant reduction in communication rates. Further investigation led us to uncover a self-jamming mechanism by which an extracellular M13 factor prevents uninfected cells from receiving the available phage signals. Model predictions of a simulated gut environment show that the novel “self-jamming” mechanism confers an advantage to both M13 and *E. coli*.

## Results

### Growth phase affects phage secretion and infection rates

To measure the rate of M13 phage secretion from growing cells, senders (TOP10_H_KanΦ, Table S1, Fig. 1a) were grown over ∼15 hours (Methods), with regular monitoring of their OD_600_ at 1h intervals (Fig. 1b), and fitted to a two-phase growth model based on Chemical Reaction Networks (Box S3, Notes S2 and S3). Briefly, growth of the bacterial cell (*C*) is represented as duplication reactions *C* + *R*_*i*_ → *C* + *C*, where *R*_*i*_ is the phase-dependent nutritional resource consumed per duplication event. The phage samples, collected at the same time-points (Fig. 1c), were quantified using the CFU assay (Methods). Phage secretion rate decreases after 12h (Fig. 1d), consistent with previous observations of reduced filamentous phage secretion rates late in the infected state^33,37^.

**Fig. 1.**
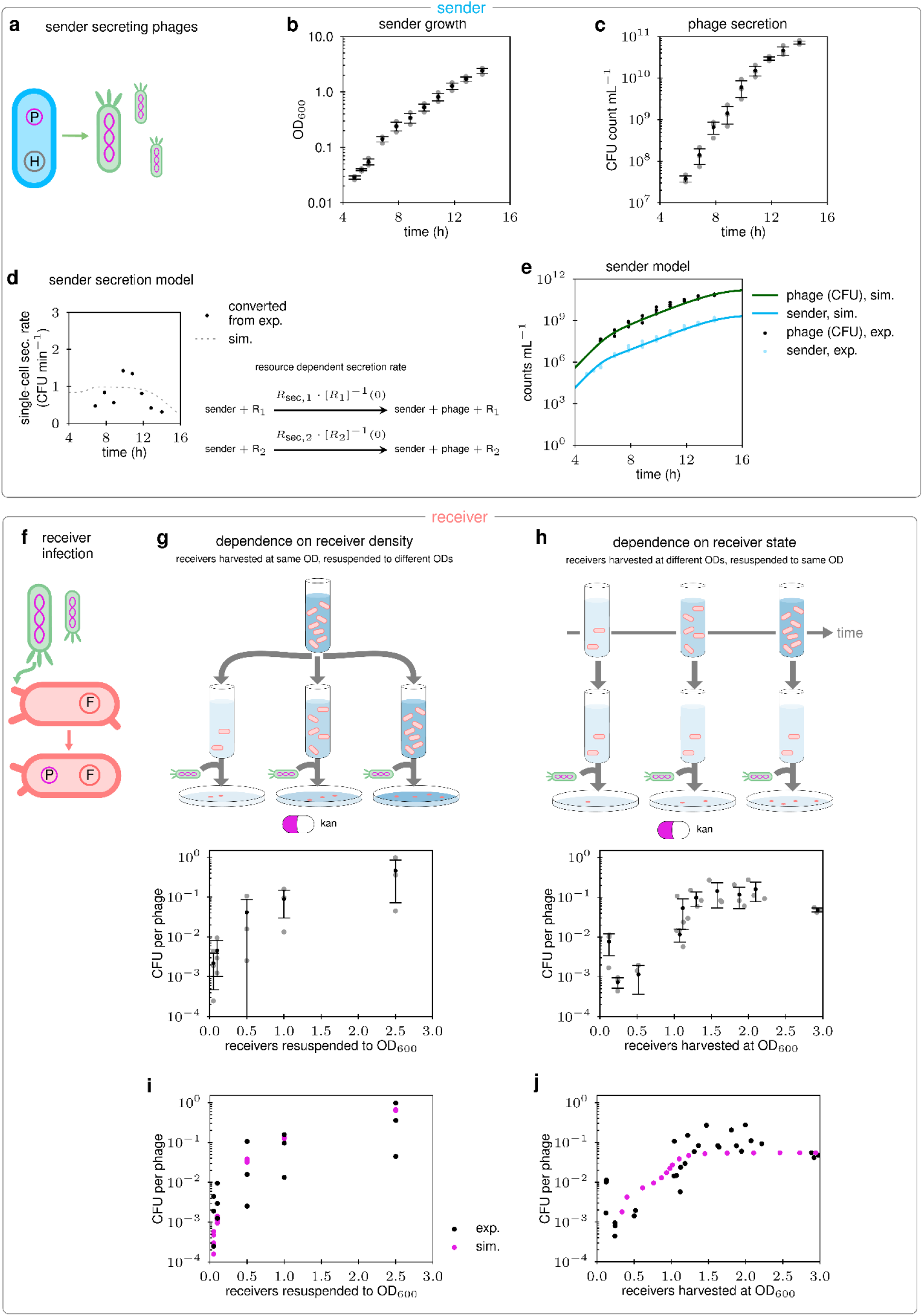
Phage secretion and infection models. **(a)** Characterisation of phage secretion by sender cells that carry a helper plasmid (H) and the phagemid (P). The latter is packaged for secretion. **(b)** Sender cells were grown in a flask (LB+antibiotics) after inoculation with overnight cultures (*n* = 3 replicates, **Methods**). OD_600_ of the senders collected over time. Data (grey), and mean ± SD (black) are indicated. **(c)** Secreted phages were collected over time and the CFU assay was used to estimate their titres (**Methods**). Data (grey), and mean ± SD (black) are indicated. **(d)** The single-cell secretion rate in CFU min^-1^ computed from (b)+(c) is shown (black) and the simulated single-cell secretion rate (grey dashed). The decrease over time was reproduced by resource dependent secretion reactions for *R*_1_and *R*_2_. **(e)** Blue: Converted sender density in CFU mL^-1^ (**Note S2**) and fitting of a two-phase growth model to the timed data. Green = secreted phage concentrations in CFU mL^-1^ and simulation of a CRN sender model that integrates the two-phase growth and phage secretion (**Note S8**). exp. = experimental data & sim. = simulation. **(f)** Characterisation of receiver cell infectability. Receivers carry an F-plasmid (F) that encodes the formation of pili, the primary phage receptors. **(g)** The dependence of infection rates on receiver densities was determined by harvesting at OD_600_ ∼1, re-dilution to different OD_600_, and subsequent CFU assay (*n* = 3 replicates, **Methods**). Data (grey), and mean ± SD (black) are indicated. **(h)** The dependence of infection rates on the receiver state was determined by harvesting receivers at different OD_600_, re-dilution to OD_600_ of 0.5, and subsequent CFU assay (*n* = 6 replicates over 2 days, **Methods**). Data (grey), and mean ± SD (black) are indicated. **(i)** Predictions of our receiver CRN model for the setup in (g) with stochastic simulations (*n* = 6 simulations per OD_600_). **(j)** Predictions of our receiver CRN model for the setup in (h).

To model secretion of phages by sender cells, we differentiated the phage counts with respect to time and normalised the secretion rates by sender cell densities (Note S4). The resulting secretion rate per cell is plotted in Fig. 1d (Note S4). To include the effect of sender physiology on secretion rates, we used a simple non-constant secretion model where the single-cell secretion rate is proportional to the cell growth rate and thus decreases with the depletion of resources. When combined with our two-phase growth model, this consists of two phage secretion reactions, one per growth reaction, with rates proportional to the growth rates. The model prediction is shown in Fig. 1d. We estimate that phage secretion rates vary from 1 phage min^-1^ cell^-1^ in the early phase to about 0.5 phage min^-1^ cell^-1^ in the late phase (Fig. 1d-e). These numbers are lower than secretion rates reported for filamentous phages between 2.5 and 21 phages min^-1^ cell^-1^ in the early phase of growth^33,36,43^, raising the possibility that our phage secretion is less efficient than for WT M13 or that the CFU assay we use to determine phage counts underestimates the true viable phage counts. Comparisons of phage counting methods show considerable variability in the reported numbers, depending on the assay method used^44,45^ (Note S5).

We next set out to investigate the effect of receiver cells’ (ER2738, Table S1, Fig. 1f) density and growth phase on the infection dynamics. Whereas in a growing culture, cell density and growth phase change simultaneously, we designed our experiments to differentiate between the two effects (Fig. 1g-h). To determine the effect of receiver densities on infection, they were grown to an OD_600_ ∼1 (late-log phase), harvested, and resuspended to five different ODs_600_, followed by infection with the same phage concentrations for a CFU assay (Fig. 1g, Methods). As expected from mass-action kinetics, we found a positive relationship between receiver cell density and the number of colonies obtained on LB agar (Fig. 1i). To determine the effect of the receivers’ growth phase on infection, they were harvested at different ODs_600_, and resuspended to an OD_600_ of 0.5. As before, the resuspended receivers were infected with the same phage concentrations for the CFU assay. We observed that the CFU counts obtained from infected receivers were ∼10-to ∼35-fold higher when the receivers were harvested in the late growth phase as compared to the early growth phase (Fig. 1h). Since the OD_600_ of the receivers was the same in these experiments, the observed differences are due to differences in infectability of the receivers at different phases of growth, likely because the number of F-pili per cell increases until the late-log phase^46^. Together, these results not only highlight the effect of receiver cell density and physiology on infection rates but also the limitations of the CFU method for accurately counting phage numbers (Note S5).

We modelled the relationship between receiver density and infection rates using mass-action kinetics (Note S8), running simulations with varying receiver densities. Using stochastic simulations (Fig. 1i, pink circles), we asked if the variance in the data (Fig. 1g) can be explained by the stochasticity of infection at small numbers. While the variance is pronounced at low OD_600_, stochastic effects in the model were found to be small overall, such that a deterministic model approximated the stochastic simulations well (Note S6). To model the higher infection rate for receivers in the late growth phase we introduced a new infectability state to the receiver model (Note S8). At the “low” infectability state, the receiver’s infection rate is only 0.4% of the “high” infectability state infection rate. We determined the rate constant of the high state infection reaction to be 3×10^−10^ mL min^-1^ from the experiments in Fig. 1g. In our model (Fig. 1j), daughter cells start at the low infectability state and transition to the high infectability state at a constant rate of 0.25 h^-1^. The distribution between low and high infectability states is determined by the initial distribution in the inoculated culture and the current OD_600_.

### Stochastic interactions influence infection dynamics of growing receiver cells

After analysing the effect of cell density and growth phase on infection of receiver cells independently (Fig. 1), we investigated how infection rates change in a growing culture as their density and growth phase change simultaneously. We set up plate reader experiments with different combinations of 4 starting receiver cell densities and 12 phage concentrations (Fig. 2a, Methods), and grew them over a duration of 20h with and without antibiotic selection (Fig. 2 and Fig. S8 to S11). In addition, we quantified unadsorbed phages from the 3^-2^ dilution at four time-points (Fig. 2c). We analysed the data to establish the killing curves due to antibiotic selection, and growth cost due to phage infection and maintenance.

**Fig. 2:**
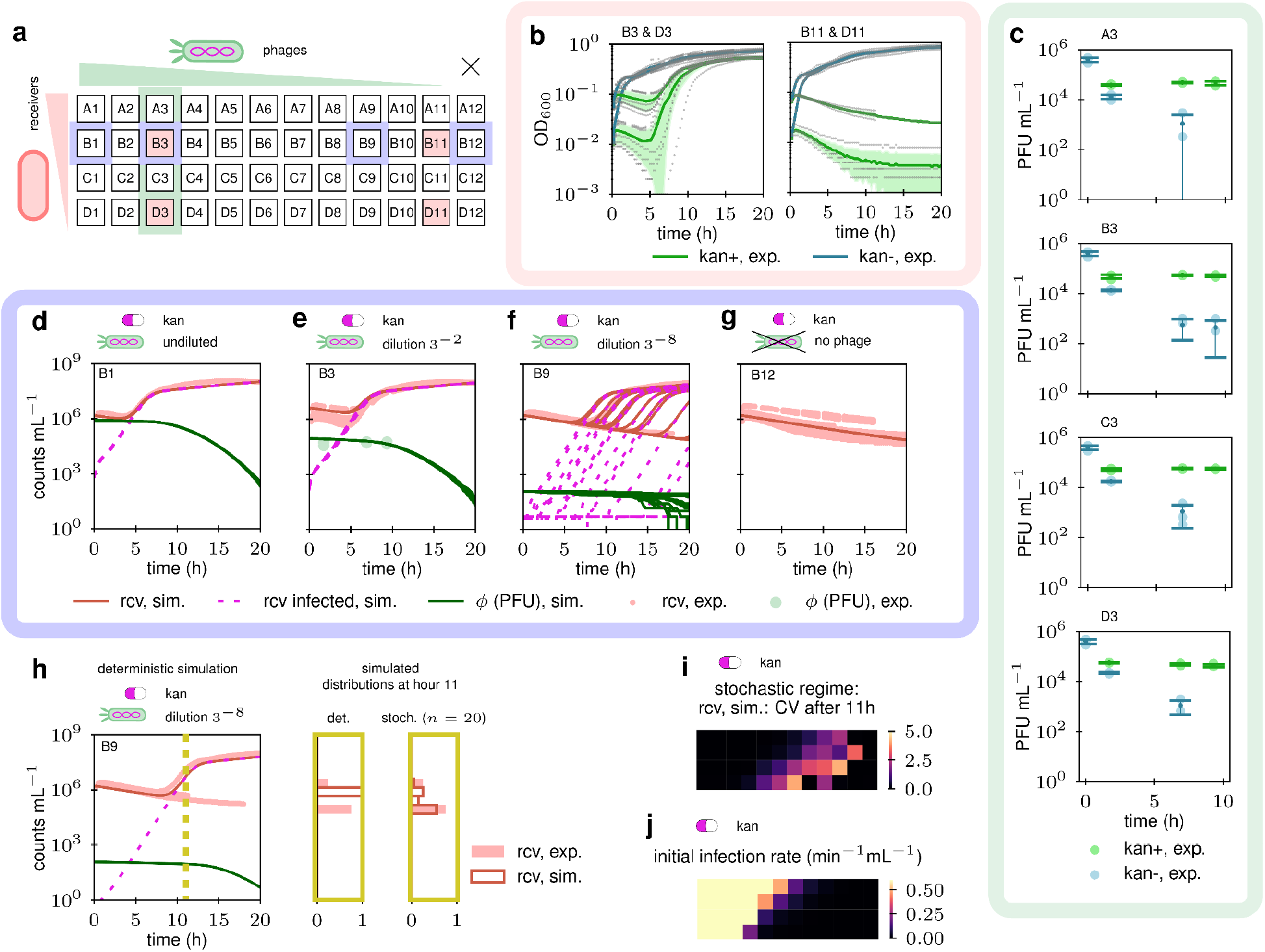
Effect of phage infection on growing receiver cells. **(a)** To receivers at 4 different cell densities, phages with 12 different concentrations were added. The OD_600_ was varied (y-axis) from 0.25 (top, undiluted) in dilution steps of ½ until 0.03125 (bottom). Added phage concentrations were varied (x-axis) from undiluted (8.16×10^5^ PFU mL^-1^, column 1), in dilution steps of ⅓ (columns 2-11). No phage was added to the rightmost column 12. Unadsorbed phage concentrations over time were measured by blue-plaque assays from column 3 (**Methods**) **(b)** OD_600_ was measured over 20h in plate reader runs +/-kanamycin (details in **Fig. S9** and **S10**). Data (grey), mean (line) and standard deviation (shaded area) are indicated. **(c)** Unadsorbed phage concentrations (PFU mL^-1^) over time for column 3 (n = 3) +/-kanamycin. Data (light circles), mean (dark circles) and SD (caps) are shown. **(d-g)** Converted receiver (rcv) cell densities (CFU mL^-1^, exp. = experimental data) and unadsorbed phage concentrations (ϕ, PFU mL^-1^) are shown, together with stochastic simulations (sim. = simulation) of our receiver CRN model (*n* = 20 runs, **Note S8**) for varying initial phage concentrations in the presence of kanamycin. For the simulations, receivers (rcv), infected receivers (rcv infected), unbound phages (ϕ, PFU mL^-1^) are shown. Plots for all wells, also in the absence of kanamycin, are shown in **Fig. S9** and **S10. (h)** Deterministic ODE simulation for the setting in (f). The different experimental outcomes are visible and reproducible with the stochastic simulation in (f) but not with the deterministic simulation in (h). The distribution of receiver densities (as fraction of the total) at hour 11 is shown in the yellow boxes for stochastic and deterministic simulations (rcv, sim), together with data from the experiments (rcv, exp). Simulations and experiments match well in the stochastic case. **(i)** Heatmap of coefficient of variation (CV) of simulated (*n* = 20 runs) receiver densities at hour 11 in the presence of kanamycin. The layout of the heatmap is the same as for (a). A diagonal region with pronounced stochastic effects is clearly visible. **(j)** Heatmap of the initial infection rates (t = 0) in the presence of kanamycin. Higher infection rates lead to likely infection and lower infection rates to likely absence of infection.

Without antibiotic (kan) selection, phage concentrations do not substantially affect the growth trajectories of cultures (Fig. 2b, S11). With antibiotic selection, receivers without phages gradually die, which is used to estimate the killing kinetics of kanamycin (Fig. 2g). However, phage infection can rescue cells from kanamycin-mediated killing, resulting in most of the kan(+) wells growing on the plate (Fig. 2d-f). When analysing the numbers of unadsorbed phages (Fig. 2c), we observe that in kan(-) wells phages continue getting adsorbed by growing cells until almost no phages are left by ∼9h. In the kan(+) wells ∼50% of the phages get absorbed within the first 2h but this number remains unchanged afterwards. As seen in multiple experimental repeats, growth in kan(+) wells is very likely in the high phage & high cell density settings and very unlikely in the low phage & low cell density settings (Fig. S9). However, in the intermediate dilutions there is substantial variability observed across experimental repeats due to the stochastic nature of contact between cells and phages (Fig. 2f).

Since the phage confers antibiotic resistance against kanamycin, we included this effect in our mathematical model. We added a kanamycin species to the CRN model, which is consumed by the receiver cells during the reaction *K* + *C* → *C*′, where *C* and *C*′ denote the cell states before and after kanamycin uptake, respectively. A receiver cell that takes up kanamycin before phage infection dies with a rate proportional to its duplication rate resulting in higher killing rates for low initial receiver densities. Such a death rate fits our data well (Fig. 2g and S9), and is in agreement with previous observations that translation-inhibiting antibiotics (like kanamycin) result in higher death rates for low initial *E. coli* densities^47^. To model the metabolic burden of phagemid maintenance after infection, we multiply each infected cell’s duplication rate by a fitted penalty factor between 0 and 1 that increases from early to late growth phase (Note S8), which is consistent with fewer phages being produced during late infection^33,37^. A reduced infection rate by factor 0.3 for cells that had taken up kanamycin prior to infection was used (Note S8). Cells transition from the early to the late infection state at a constant rate of 1/3 h^-1^ (Note S8). The model allows us to generate timed simulation traces involving infection and antibiotic killing-resistance kinetics. We used deterministic and stochastic simulations to predict OD_600_ trajectories across several experiments over 4 plate runs (Fig. 2d-g, S9, and S10).

While in the kan(-) plate reader experiments, deterministic simulations approximate stochastic simulations well (Fig. S10), this is not the case in the kan(+) experiments. The stochastic simulations (Fig. 2f) show that infected receivers (dashed) are low in number over a long period of time, and when they do grow there are large variations in the starting time depending on the initial receiver density. This results in large variations of the receiver densities after 5 hours, a behaviour that the stochastic simulations can reproduce (Fig. 2f), but the deterministic simulations cannot (Fig. 2h). The heatmap in Fig. 2i shows where stochastic simulations lead to large variations in receiver growth by reporting the coefficient of variation (CV) for receiver counts at hour 11. The darker top-left and bottom-right corners of the heatmap correspond to the setups where infection is either likely to occur or likely not to occur, following a deterministic pattern. The highest CV is seen along the diagonal where the lower numbers of phages or receivers result in high variability in infection outcomes. This can also be seen by considering the initial infection rates (Fig. 2j). The stochastic region corresponds to an expected first infection within the first few minutes, after kanamycin-mediated killing starts to significantly decrease the uninfected *E. coli* counts.

### Resource-sharing and antibiotic selection during communication between sender and receiver cells

We next studied the dynamics of intercellular communication between sender and receiver cells growing in a co-culture. To assess the communication between them via phages (pSB1K3_M13ps_LacZα_gIII), senders (TOP10_HΔgIII_gIII-KanΦ) and receivers (ER2738) were incubated without antibiotic selection for 1h (Fig. 3, Methods), following which they were grown in 4 experimental sets over 15h: one without antibiotics, and three with antibiotics to select for senders only (gentamycin), receivers only (tetracycline), or infected receivers only (kanamycin+tetracycline) (Fig. 3a-e and S18). To simulate these results, the sender secretion and receiver infection models were combined, and effects of gentamycin and tetracycline selection added (Note S8). Experimental results and deterministic ODE simulations are shown in Fig. 3f-i, and S17 to S22. Stochastic simulations (not shown) showed no apparent difference to the deterministic simulations.

**Fig. 3:**
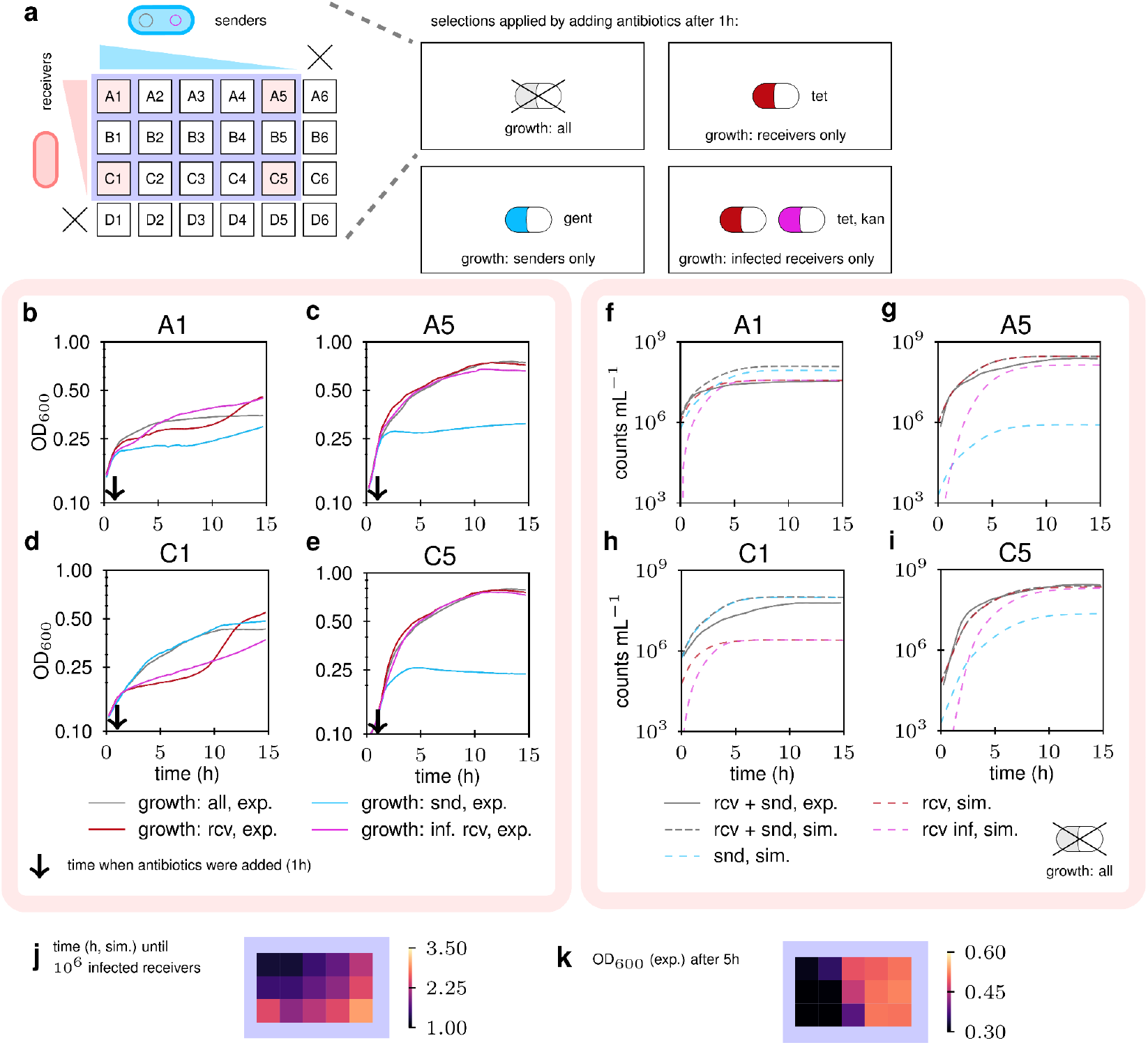
Communication from senders to receivers. **(a)** Senders (snd) and receivers (rcv) were mixed in different combinations of their starting ODs and grown for 15h (in four sets), with different antibiotic selections applied to three sets after 1h (**Methods**). For each set, initial sender (x-axis) and receiver densities (y-axis) were varied, starting from the undiluted (OD_600_ = 0.136), in dilution steps of ½, with the rightmost/bottommost column/row having no sender/receiver. The four sets correspond to: no antibiotics added, tetracycline to select for receivers only, gentamycin to select for senders only, and tetracycline+kanamycin to select for infected receivers only. **(b-e)** OD_600_ over 15h was monitored in a plate reader run (n = 1). Four representative wells are shown with the time-point of antibiotic addition indicated by an arrow. Each well shows curves for all four sets of antibiotic selection. Plots for all wells are shown in **Fig. S18. (f-i)** Converted cell densities (rcv+snd, exp.) are shown with deterministic simulations of the combined sender and receiver CRN model (**Note S8**) for the settings (b) to (e). For the simulations, total cell (rcv+snd, sim.), sender (snd, sim.), receiver (rcv, sim.), and infected receivers (rcv inf, sim.) densities are shown. As before, plots for all wells are shown in **Fig. S17** and **S19** to **S22. (j)** Heatmap for the purple area of (a) with simulated time (h) until the density of infected receivers reaches 10^6^ mL^-1^ in the absence of antibiotics. The earliest time is reached with high densities of both senders and receivers (top-left). **(k)** Heatmap for the purple area of (a) with experimentally measured OD_600_ after 5h in the absence of antibiotics. Higher ODs are obtained with low initial sender densities (right).

In the absence of antibiotic selection, final ODs are lower when starting sender densities are high (Fig. 3b&d) due to the senders growing slower than the receivers while consuming the same amount of resources (Fig. 3a-e, S18). This is confirmed by the model where senders dominate the final numbers in wells A1 and C1 (Fig. 3f&h), but receivers dominate the numbers in wells A5 and C5 (Fig. 3g&i). A similar effect is also observed when selecting for receivers, where higher starting sender densities lead to lower final ODs due to early resource competition before the senders are killed by the antibiotic (Fig. 3b-e), resulting in apparent biphasic growth at high starting sender densities (Fig. 3b&d). The model simulations confirm the overtaking of the dying senders by the receivers (Fig. S20). The reverse is observed when selecting for senders: final ODs are higher in wells with lower starting receiver densities (Fig. 3b&d), as confirmed by the model (Fig. S21). When selecting for infected receivers, the starting high-sender settings (Fig. 3b&d) show differences in growth due to the killing of senders and uninfected receivers, followed by the subsequent growth of infected receivers. These complex dynamics result in the observed biphasic growth curves that model simulations agree with (Fig. S22).

Taken together, these data indicate that in the absence of selection most phage transmission is horizontal while in the presence of selection most transmission is vertical. The average transmission time between senders and receivers at the end of the simulations (Fig. 3f-i) is 11–1334 min in the absence of selection and 11–21 min in its presence. The simulations explain the experimental data well (Fig. 3f-i, S18, S19 to S22), with a heatmap of the time taken to reach 10^6^ infected receivers showing that starting high densities of both senders and receivers lead to faster infection rates (Fig. 3j). However, higher final infected receiver densities are achieved when starting with lower sender densities due to the lower starting competition facilitating later post-infection growth (Fig. 3k). Interestingly, the stochastic effects seen earlier due to low phage and cell numbers (Fig. 2f) are not seen when using sender cells for phage delivery to comparable densities of receivers, offering a possible explanation for why delivery of sender cells rather than isolated phages shows better phage titres in certain gut applications^48^.

### Conditioned media inhibits phage infections

To understand how the infection dynamics would be affected by the state of the media in long-term experiments, e.g., because of spent media reducing the growth differences between uninfected and infected cells, we collected conditioned media from senders (TOP10_H_KanΦ, containing Helper+KanΦ, Table S1) and empty-vector control cells (TOP10_Gx_Kx, Table S1), and removed cells with a 0.22 µm filter (Methods). The filtrate and phages with a different antibiotic resistance marker (AmpΦ) were then added to receiver cells (ER2738) prior to monitoring their growth under ampicillin selection (Fig. 4a). In contrast to fresh LB media and empty-vector conditioned media, a delayed growth was observed in the sender conditioned media. To rule out infection competition by phages secreted by the senders (KanΦ), they were removed from the sender conditioned media by passing through a 50kDa filter (fraction B, Fig. 4c, Methods). Infection assays with fraction B of the sender conditioned media also showed delayed growth, although a little earlier than fraction A (Fig. 4b).

**Fig. 4:**
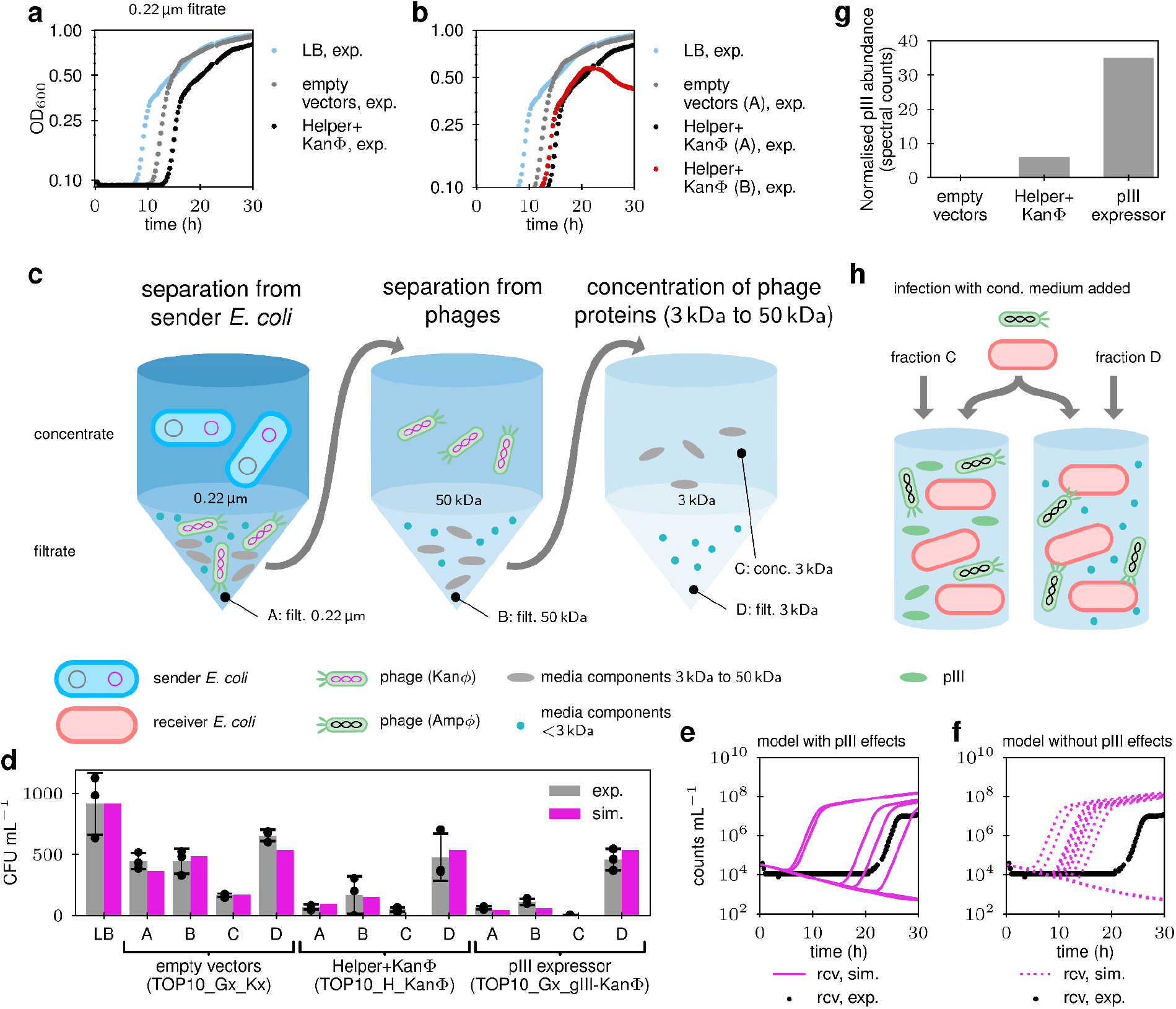
Extracellular secretion of M13 pIII and its inhibitory effect on infection. **(a)** Conditioned media from sender cells (TOP10_H_KanΦ, carrying Helper+KanΦ) and cells carrying empty vectors (TOP10_Gx_Kx) were filtered to remove the cells. The two conditioned media (48% concentration) and an LB control were then mixed with AmpΦ phages, incubated with receiver cells for infection, and the infection reactions transferred to fresh for growth over 30h (**Methods**). Receiver growth (OD_600_) in LB+ampicillin is shown (n = 1). Plate reader runs of growth from other infections in conditioned media, and their dilutions, are shown in **Fig. S23**, +/-ampicillin. **(b)** Data from (a), but in addition with sender conditioned media (fraction B) after passing through a 50kDa filter to remove KanΦ phages (48% concentration) (n = 1, see **Methods** and **Fig. S23** for additional growth data). **(c)** Preparation of filtrates (A, B, and D) and concentrate (C) by passing conditioned media through 0.22 µm, 50kDa, and 3kDa filters (**Methods**). **(d)** Subsequent infection assays of receiver cells with AmpΦ phages were carried out in an LB control and fractions A to D of three conditioned media (n = 3, see **Methods**). Data (black dots), mean (grey bars), and SD (black caps) are shown for LB and conditioned media fractions A to D from three cell cultures (Table S1; volumes of the fractions in **Table S3**). Simulations of our mathematical model for inhibition of infection by pIII in the conditioned medium are shown in pink (see **Note S7** for the model). **(e-f)** Receiver growth over 30h (black) with ampicillin selection (details in **Fig. S23**), following infection in fraction B of the conditioned media (80% concentration) from sender cells. Stochastic simulations (n = 16) of receiver densities during growth, obtained from our CRN model with infection inhibition by pIII (**Note S8**), are shown (rcv, sim.) in (e) for the full model, and in (f) when disregarding infection inhibition by pIII. The model in (f) incorrectly predicts too early growth, or no growth, of receiver cell densities. **(g)** pIII abundance in conditioned media from the three cell cultures, as determined by secretome analysis (LC-MS/MS, **Methods**). **(h)** Schematics of infection assays of receiver cells with AmpΦ phages when adding fraction C versus fraction D of the sender/ pIII expressor conditioned media.

Intrigued by the above results, we next performed a CFU assay to quantify the number of phages when the infection is carried out in different conditioned media fractions (Fig. 4c). We find that fractions A, B, and C of conditioned media from the sender cells (TOP10_H_KanΦ) reduce the number of infected colonies by 92%, 82%, and 94% compared to the LB control, respectively, while infected colonies in fraction D are only reduced by 48% (Fig. 4d). This indicates that a component present in fractions A, B, and C of the sender medium, but absent in fraction D, is responsible for the infection inhibition. It follows that this component is between 3 and 50 kDa in size (all known M13 proteins are between 3kDa and 47kDa) and is uniquely present in the conditioned media from the senders, but not from the parental strain. Taken together with control data, the reason for reduced infection efficiency in the sender conditioned media appears to be a sender-specific molecule in its secretome (fraction A, Fig. 4d) that reduces the percentage of successfully infecting phages in the population by ∼6-fold (from 5% to 0.8%).

We wondered if M13 encodes an extracellular factor that causes infection immunity against itself. Adding purified M13 pIII protein to the extracellular medium is known to inhibit conjugation in *E. coli* by blocking the F-pilus^49^. Given that the F-pilus is also the surface receptor for M13 entry^38,50^, we hypothesised that the unknown factor causing extracellular infection immunity in our experiments is the minor coat protein pIII. To test this hypothesis, we constructed a strain that expresses pIII in *E. coli* TOP10 cells (TOP10_Gx_gIII-KanΦ), but no other M13 phage protein, and used it to make conditioned media. Indeed the pIII expressor conditioned media showed a similar infection inhibition effect as the sender conditioned media, in both the plate reader (Fig. S23) and CFU assay experiments (Fig. 4d). To directly test if the pIII protein is present in the conditioned media, we sent fraction C from all the three strains (parental, sender, and pIII expressor) for mass-spectrometric analysis (Methods, Fig. 4g). The results show that pIII is present in the conditioned media prepared from the sender and the pIII expressor cells, but not from the control cells, confirming pIII as the extracellular factor responsible for M13 infection immunity. This also explains why fraction D does not inhibit infection but fraction C does (Fig. 4h).

We modelled the CFU assay data (Fig. 4d) with a first-order kinetics infection model (Note S6). The infection rate is modulated by viscosity factors (*visc*, for A to D) and pIII concentration factors (*pIII*) that reduce the density of susceptible receivers to *R*_*S*_, with exponential dependence on *pIII* (first-order kinetics). The predicted CFU count (*CFU*) is then obtained as 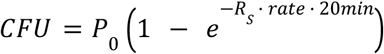 with *rate* being inversely proportional to the viscosity (Stokes-Einstein) and *P*_0_ being the initial concentration of phages. After including these factors, simulations from the model explain the data well (Fig. 4d). The infectability of cells (Methods) in fraction A from the sender conditioned media was significantly lower than that from the control (p = 0.005). However, no significant difference was seen between fraction A from the sender and the pIII expressor (p = 0.640). The calculated infection rates are reduced by 70% when going from the control to the sender conditioned media. To evaluate the models with and without pIII inhibition, we compared the data from the conditioned media fraction A against both models (Fig. 4d, S25). We found a significant difference between the data and the model when the pIII effect was absent (p = 0.015 for control, 0.091 for sender, 0.004 for pIII expressor) but not when it was present (p = 0.168, 0.172, and 0.185, respectively).

A dynamic model was obtained from the receiver model (Note S8), comprising a pIII jamming reaction for unjammed receivers (mass-action kinetics), and an unjamming reaction (first-order kinetics) for jammed receivers. Unjamming occurs upon cell duplication in one of the two daughter cells. The best fittings (Fig. 4e) were obtained with a single daughter cell becoming unjammed upon growth with resource *R*_2_. A control model where pIII secretion is removed does not fit the experimental data (Fig. 4f), predicting a too early growth of infected receivers.

### Extracellular infection inhibition is advantageous for both *E. coli* and M13

Bacteria and phages are perpetually engaged in an arms race, where they try to gain fitness advantage over each other^51^. Having discovered that the M13 minor coat protein pIII secreted by infected cells blocks the entry of M13 phage into uninfected *E. coli* cells, we next used the CRN model to ask whether this mechanism benefits *E. coli* or M13 in the gut environment. We simulated the gut environment as a continuous culture system where resources *R*_1_ and *R*_2_ flow in, and bacteria (*E. coli* and a competitor), phage M13, secreted pIII, and the resources flow out. The in- and matched out-volume basal flow-rate (R) was set around 5 mL min^-1^, as found in literature^52–54^. *E. coli* and competitor cells flow at a rate lower by a factor (f) due to adhesion to the intestinal mucosa^40^ (Fig. 5a). The resources are consumed by the *E. coli* to grow and, if infected, the *E. coli* produce phages and secrete pIII. The competitor cells feed on the same resources *R*_1_ and *R*_2_ and have a growth rate slightly higher (+1%) than uninfected *E. coli*. Infection of *E. coli* leads to a reduced growth rate, resulting in substantial disadvantage against the competitor cells.

**Fig. 5:**
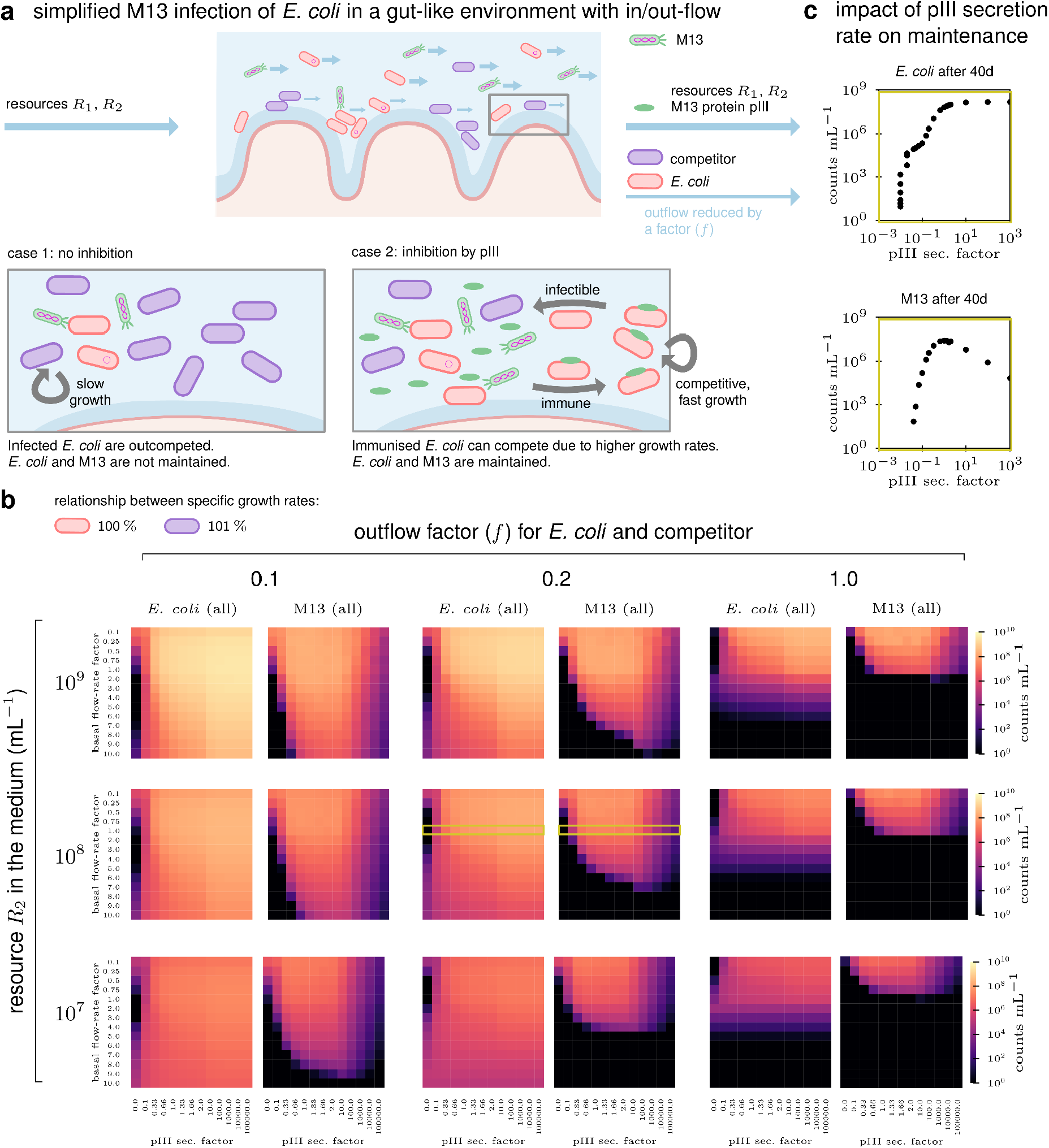
Simulation of long-term infection with and without self-jamming of M13 phages. **(a)** A simple CRN model (**Note S8**) for M13 in a gut-like environment that combines secretion and infection capabilities from the sender and receiver models into a single *E. coli* host (red), feeding on resources *R*_1_ and *R*_2_. A competitor cell (blue) feeds on the same resources. The gut-like environment is modelled by an in-flow of resources and a matched out-flow of resources, M13, protein pIII, and *E. coli* and competitor cells. The *E. coli* and the competitor flow out at a rate reduced by the outflow factor *f* due to adhesion to the gut mucosa. **(b)** Simulations for the case where the competitor cell has a growth rate 1% higher than the uninfected *E. coli* (also see **Fig. S30 & S31**). Counts mL^-1^ of all *E. coli* (infected and uninfected) and all M13 phages (inside and outside cells) after 40d are shown in each heatmap tile for varying relative basal flow-rates (rows = inner y-axis, factor of 1 = 5 mL min^-1^) and varying pIII secretion rates (columns = inner x-axis). The tile column with pIII secretion factor 0 corresponds to the case without infection inhibition via pIII. The heatmap tiles in turn are organised according to varying resource *R*_2_ concentrations (outer y-axis) and varying outflow factors *f* (outer x-axis). The initial density of *E. coli* (with a high infectability fraction of 10%) and the competitor cells is 10^3^ mL^-1^ (changes up to 10^5^ mL^-1^ showed no visible difference). **(c)** Counts of *E. coli* and M13 after 40d for varying secretion rate factors (resource *R*_2_ = 10^8^ mL^-1^, *f* = 0.2, basal flow rate factor = 1.0). The corresponding parameter range in (b) is marked in yellow.

Model simulations over a prolonged time (Fig. 5b, S26 to S28) show that the infection inhibition mechanism due to pIII secretion is beneficial to both *E. coli* and M13. When the *E. coli* are most sticky (f = 0.1), the resource concentration is high (R_2_ = 10^9^ mL^-1^), and the basal flow-rates are medium (factor 2-6) (Fig. 5b), both *E. coli* (uninfected+infected) and M13 (inside+outside cells) numbers are 4-6 orders of magnitude higher than would be expected without pIII secretion. In these settings, and for a given secreted pIII concentration, as the basal flow-rate increases the *E. coli* numbers reduce slightly but the M13 numbers reduce drastically. This indicates that at higher basal flow-rates, the relative advantage of pIII secretion for M13 is much higher than that for *E. coli*. When the *E. coli* are less sticky (f = 1), the pIII secretion advantage still exists, but quickly diminishes with increasing basal flow-rate. The pIII secretion rate was varied from 0.1 to 10^5^ times the nominal rate and found to benefit M13 over a large range, except at an unrealistically high rate where it leads to washout (Fig. 5c).

Fig. S26 to S28 show how pIII secretion ensures higher *E. coli* and M13 numbers in the presence of competitor cells by maintaining a sub-population of jammed, and hence uninfectable, fast-growing receivers. *E. coli* inhibited by pIII have growth rates close to the competitor’s and constantly replenish infectable receivers, ensuring that M13 are not washed out. Fig. S29 shows a control simulation without pIII secretion. We also varied the competitive pressure on the *E. coli* by setting the growth rate of the competitors equal to uninfected *E. coli* (Fig. S30) and 5% higher (Fig. S31). We found that the pIII advantage is more pronounced against a faster-growing competitor: compare, e.g., the bottom-left tile where the range of flow rates for which pIII secretion is advantageous is larger.

## Discussion

Here, we have developed an intercellular communication system that uses the M13 phage machinery to package ssDNA for transmission from sender to receiver cells. Owing to the high programmability of DNA^55,56^, the system has the potential to generate a large number of orthogonal phage signals to dramatically expand the communication alphabet for cell-to-cell communication^57^. An expanded alphabet in turn would increase the communication capability in multicellular circuits that are currently highly constrained by the small number of orthogonal small-molecule based signals^58,59^. With the large M13 packing capacity of 40 kb DNA^60^, phage signals are amenable to delivery of a wide range of genetic programs, from fluorescence and antibiotic markers^57^ to metabolic pathways^17^, toxic peptides^12^ and sequence-specific antimicrobials^14^. Phage signals also have a high speed of intercellular transmission, with the average time of sender-to-receiver communication being as low as 11 min in our simulations, even without selection (Fig. 3g). The time until 10^6^ infected receivers ranges from 1.25h to 3.5h in our setup (Fig. 3j). For a phagemid size of 5.6 kb this represents an average bitrate of 2.5 bit per second (1.25 bases per second). While significantly lower than the first modems (∼100 bit per second^61^), it is not much lower than the intracellular information transmission via transcription as seen in *E. coli*^*62*^.

As a well-studied filamentous phage, the kinetics of the natural M13 infection cycle have been known and modelled for many years^33,34,36–38,63^. However, detailed population-level models for a wide range of growth conditions were missing. In this work, we develop such population-level models for engineered phages where infection and packaging functions have been separated into different DNA molecules. Inspired by the communication primitives in networked computer systems^64,65^, for a comprehensive and realistic description of the signalling process, our quantitative CRN models cover a wide range of growth conditions and include several features of phage communication not previously seen together: maintenance burden of the communication machinery and the received signal, cell density and growth-phase dependent differences in signalling dynamics, antibiotic selection effects, as well as stochasticity of events. The stochastic effects in our models are key to explaining infections at low receiver and phage concentrations. It was previously hypothesised that the stochastic nature of phage-bacterial interactions due to spatial heterogeneity in the gut is responsible for the poor efficacy of oral phage therapy^66^. Our results suggest that such stochasticity may be overcome by using sender cells instead of isolated phages for phage delivery.

In our attempt to engineer the phage-mediated communication system, we have also obtained unexpected insights into it. We discovered that conditioned media from phage senders inhibits phage infection. Although superinfection immunity due to intracellular phage proteins has been reported in many systems: sieB of *Salmonella* phage P22 and *E. coli* phage lambda^67^, pIII of *E. coli* phage M13^68,69^ (see Fig. S24), and PfsE of *Pseudomonas* phage Pf^70^, to the best of our knowledge, this is the first reported example of extracellular release of a phage protein to modulate immunity against the phage itself. Although mechanistically dissimilar, it is reminiscent of immune evasion by respiratory syncytial virus mediated by its extracellular soluble G protein^71,72^. Our data show that the pIII is present in the extracellular medium and protects uninfected *E. coli* against M13 infection, settling an open question about an unknown “diffusible factor” that was speculated upon >40 years ago^35^.

Bacteria and phages are perpetually engaged in a coevolutionary arms race^73,74^. In obligate parasitic relationships like these, the molecular mechanisms of one-upmanship get so intertwined during long-term association that the fitness advantage of the bacterium and the phage may get mutually aligned^75^. This makes it harder to assess the success of the phage parasite or the host bacterium separately, leading to suggestions that a “virocell” paradigm that sees the infected host as a distinct organism is a more appropriate way of assessment^76^. Here, we use three different measures (free phages, infected cells, free phages+infected cells), and found that our results hold across all measures (Fig. 5b, S26-S31). The self-jamming mechanism we report protects a fraction of uninfected bacteria from M13 infection, allowing them to multiply at higher rates. In turn, this makes a fresh crop of bacteria available to M13 for future infections. This resembles the host-farming practices of the *Salmonella* phage P22 where a population of susceptible hosts is generated following asymmetric cell division of an infected host^77^. In addition to informing future phage therapy applications, our findings can have key implications for technologies like phage display and phage-assisted continuous evolution.

## Methods

### Bacterial strains, growth conditions, and cloning

All *Escherichia coli* strains were grown at 37 °C in LB media (liquid with shaking at 200 rpm, or solid LBA plates with 1.5% w/v agar) supplemented with the appropriate antibiotics at the following concentrations (unless otherwise indicated): kanamycin (kan 30 µg mL^-1^), ampicillin (Amp 100 µg mL^-1^), gentamycin (gent 10 µg mL^-1^), tetracycline (tet 10 µg mL^-1^), and streptomycin (Str 50 µg mL^-1^); concentrations were halved when using dual antibiotic selection. Strains and antibiotics used are listed in Table S1 (see also: Note S1). To calculate the growth rates of strains, cells were diluted 100x from overnight cultures and re-grown in a 96-well plate (200 µL per well) in a plate-reader (Biotek Synergy HTX) until they reached an OD_600_ of 0.2 to 0.3, following which they were diluted again by 40x into a new plate. Cultures in the second plate were grown overnight, recording their OD_600_ at 15 min intervals. The data generated was used to calculate the Specific Growth Rates (Fig. S7).

Cloning was performed by Golden Gate Assembly of PCR-amplified DNA fragments using NEB enzymes: Q5 DNA polymerase (#M0492), BsaI-HFv2 (#R3733), and T4 DNA ligase (#M0202M). *E. coli* strains Dh5α and TOP10 were used for cloning. All plasmids constructed were verified by Sanger sequencing. Maps of plasmids used in this study are presented in Fig. S2 and the sequences listed in Table S1.

### Absorbance-cell density relationship

The relationships between absorbance (OD_600_) and cell density for the two strains (Fig. S5 a&b, S6) were determined by diluting by 1000x an overnight culture and re-growing at 37 °C in fresh LB media. Aliquots were taken from the growing cultures at several time-points and their absorbance (OD_600_, spectrophotometer) recorded, cooled down on ice for ∼30 min, following which they were serially diluted, 10^−4^ to 10^−7^ dilutions were spread on LBA plates, and the colony forming units (CFU) obtained were counted the next day. In a separate experiment (Fig. S4), the relationship between the absorbance measured using a spectrophotometer (UVisco V-1100D) and a plate-reader (Biotek Synergy HTX) was determined, and the relationship with cell density accordingly re-calculated (S5 c&d).

### Phage secretion

To determine the effect of cell physiology on phage secretion, three independent cultures of sender cells (TOP10_H_KanΦ) were grown in shaker flasks to different cell densities (spectrophotometer OD_600_ ∼ 0.05, 0.1, 0.2, 0.3, 0.4, 0.6, 1.0, 1.5, 2) from 1000x diluted overnight cultures, 1 mL each of the cultures was spun down at 4000x g, the supernatant filtered using a 0.22 µm filter, and the filtrate (phage prep) saved at 4 °C. Subsequently, the receiver strain (ER2738) was grown to OD_600_ ∼1 and used to perform the CFU assay to determine the titres in the phage preps obtained above. Phage preps from other sender cells (TOP10_H_AmpΦ) were similarly prepared from an overnight grown culture and quantified using the CFU assay. Preps from sender cells that produce plaque-forming phages (TOP10_H_gIII-KanΦ, TOP10_HΔgIII_gIII-KanΦ) were quantified using the CFU assay (receiver: ER2738 or ER2738_HΔgIII) and/or the PFU assay (receiver: ER2738_HΔgIII).

### Phage infection, and CFU/ PFU assays

Phage titres were quantified using either a CFU or a PFU assay. Both assays involve mixing a phage dilution with *E. coli* receiver cells (containing an F-plasmid) followed by: either (1) a period of incubation and then spreading onto a selective LBA plate (CFU assay), or (2) immediate mixing with soft agar and pouring on a solid LBA plate (PFU assay).

For the CFU assay, receiver cells grown overnight were diluted 1000x and re-grown at 37 °C in LB (with appropriate antibiotics) until they reached a spectrophotometer OD_600_ between 1 and 1.5. Cells were chilled on ice for at least 30 min, and then 90 µL aliquoted into eppendorf tubes. The tubes were moved to room temperature (RT) for 5 min before adding phages to the cells. 10 µL of different phage dilutions (10^−1^ to 10^−7^) were mixed with the receiver cells and incubated at RT for 20 min. Thereafter, the mixtures were plated on LBA plates with the appropriate antibiotic concentration. Colonies on the LBA plates were counted the next day after incubation at 37 °C for ∼16h. Counts from plates between 30 and 300 colonies were used to determine the mL^-1^ titres of the phage preps according to the formula: CFU count / (phage dilution * phage volume used in mL). Since the CFU assay relies on the ability of the phage to confer antibiotic resistance to an infected receiver cell, it can be used to quantify any phage in this study using any of the receiver cells (Table S1). For the experiments in Fig. 1 (g&i), the receivers were harvested at OD_600_ ∼1 and cooled, and their density was later adjusted to different OD_600_ values (0.05, 0.1, 0.5, 1.0 and 2.5). For the experiments in Fig. 1 (h&j), the receivers were harvested at the different OD_600_ indicated and cooled, and their density later adjusted to OD_600_ of 0.5.

For the PFU assay, receiver cells (ER2738_HΔgIII) grown overnight were diluted 1000x and re-grown at 37 °C in LB (tet+gent) until they reached a spectrophotometer OD_600_ between 1 and 1.5. Cells were chilled on ice for at least 30 min, and then 90 µL aliquoted into eppendorf tubes. The tubes were moved to RT shortly before mixing 10 µL of different phage dilutions (10^−1^ to 10^−7^) with the receiver cells, and then adding the mix to 10 mL of soft LBA (0.75% w/v agar), previously aliquoted into a 15 mL tube and kept molten at 50 °C. The phage+receiver mix in the soft agar was immediately poured onto a solid plate with 20 mL hard LBA (1.5% w/v agar) with 0.2 mM IPTG and 40 µg mL^-1^ X-gal, and after the soft LBA had solidified the plate was incubated at 37 °C for 16-24h. Plaques of the non-lytic M13 phage are turbid/ diffused, usually making them harder to see. IPTG and X-gal in the hard LBA colours the plaques blue (LacZω in the F-plasmid is complemented by the LacZα in the phagemid), making them easier to visualise (Fig. S15). Counts from plates between 30 and 300 plaques were used to determine the mL^-1^ titres of the phage preps according to the formula: PFU count / (phage dilution * phage volume used in mL). Since the PFU assay relies on the phage infecting a receiver cell and getting re-secreted to infect the neighbouring receiver cells in the soft agar, it can only be used to quantify phages carrying gIII (pSB1K3_M13ps_LacZα_gIII) that infect receivers containing an M13 Helper plasmid without gIII (ER2738_HΔgIII) (see Table S1).

To determine the relationship between CFU and PFU titres obtained from the same phage prep (pSB1K3_M13ps_LacZα_gIII), three independent colonies of receiver cells (ER2738_HΔgIII) were grown to a spectrophotometer OD_600_ of 1-1.5 in LB (tet+gent) and used for both the CFU and the PFU assays (see Note S5 and Fig. S16).

### Monitoring of growth and infection in a plate-reader

For the infection plate run experiments of Fig. 2, overnight cultures of receiver cells (ER2738) were diluted 1000x and re-grown to a spectrophotometer OD_600_ of 0.6-0.7, following which they were cooled on ice for ∼30 min and their OD_600_ re-adjusted to different densities (0.25, 0.125, 0.0625. 0.03125) while still on ice. Several serial dilutions (3^N^-fold, N = 0 to 10) of the phage prep (pSB1K3_M13ps_LacZα_gIII, undiluted concentration of 36.2×10^5^ PFU mL^-1^), and a no-phage control, were prepared and 45 µL aliquoted into a 96-well plate at RT. 150 µL of the different receiver dilutions were added to the plate and incubated at RT for 20 min. Next, 5 µL of LB was added to each well, without or with kanamycin (end concentration 30 µg mL^-1^), and the plate grown overnight while recording OD_600_ at 15 min intervals. The above experiments were repeated four times, each with a different set of phage dilutions added to the plate: (a) 3x technical replicates of dilutions 3^8^-3^10^ and the no-phage control (continuous), (b) 3x technical replicates of dilutions 3^1^-3^3^ and the no-phage control (continuous), (c) 3x technical replicates of dilutions 3^1^-3^3^ and the no-phage control (discontinuous), and (d) single replicate of dilutions 3^0^-3^10^ and the no-phage control (continuous). All data from the four repeats were combined before plotting/ model-fitting. In the discontinuous run, the plate was paused at several time-points (0, 2, 6, and 10h) to draw a 3 µL sample from each well of the third column (phage dilution 3^2^), which was added to 200 µL of LB (gent) to kill all cells and later used to quantify by PFU assay the unadsorbed phages in the well (Fig. 2c). The pauses for phage sampling resulted in an average gap of ∼45 min between plate reader measurements before and after the pause, resulting in the plotted time-points in Fig. 2c being accordingly delayed.

For the communication plate run experiments of Fig. 3, an overnight culture of receiver cells (ER2738) was diluted 10000x and re-grown to a spectrophotometer OD_600_ of 0.4, following which it was cooled on ice for ∼30 min and several OD_600_ dilutions made (0.136, 0.068, 0.034, and 0.0) while still on ice. Overnight culture of senders (TOP10_H_gIII-KanΦ) was diluted 100x and re-grown to a spectrophotometer OD_600_ of 0.2, following which it was cooled on ice for ∼30 min and several OD_600_ dilutions made (0.136, 0.068, 0.034, 0.017, 0.0085, and 0.0) while still on ice. 90 µL of receiver cell dilutions were added per well to a 96-well plate in quadruplet (see Fig. 3a for the organisation of the four sets), followed by 90 µL of the sender cell dilutions also in quadruplet. The plate was run at 37 ºC for 1h, following which 20 µL of LB with the appropriate antibiotics (5x concentrated, to achieve the standard end-concentration) was added to each well, and the plate was grown overnight while recording OD_600_ at 15 min intervals.

### Phage infections in the conditioned media fractions

Cultures of six strains (see Fig. S23, Table S1) were grown overnight in LB (gent+kan), spun down at 4000x g, the supernatant filtered using a 0.22 µm filter (Millipore SLGP033RS) to remove any cells, and the filtrate saved at 4 ºC (fraction A, conditioned media). Of these strains, since TOP10_H_KanΦ is the only one expected to produce phages, conditioned media (fraction A) from it was also filtered using a 50 kDa filter (Millipore UFC905024) to obtain phage-free conditioned media (fraction B). Six concentrations (0% = fresh LB control, 16%, 32%, 48%, 64%, and 80%) of these conditioned media were used as background to run plate-reader infection experiments (as above) using an AmpΦ phage (pSEVA19_M13ps). Briefly, 160 µL of the six media dilutions prepared earlier from each of the seven media types (fraction A from six different strains as well as fraction B from TOP10_H_KanΦ) were added to a 96-well plate, followed by 20 µL of the AmpΦ phage (undiluted concentration 1.85×10^4^ CFU mL^-1^). Finally, 20 µL of ER2738 receivers (grown to a mid-log phase and resuspended to OD_600_ of 2.6) were added to each well (effective OD_600_ of 0.26) and incubated at RT for 20 min. Following the incubation, 10 µL of the above infection reactions were added to a new 96-well plate containing fresh LB (without and with 100 µg mL^-1^ ampicillin), and the plate grown overnight while recording OD_600_ at 15 min intervals. The data are plotted in Fig. 4a-b, 4e-f, and S23. 200 mL cultures of a subset of the strains used above (TOP10_H_KanΦ, TOP10_Gx_Kx, and TOP10_Gx_gIII-KanΦ, Table S1) were grown overnight in LB (gent+kan). Each culture was spun down at 4000x g, the supernatant filtered using a 0.22 µm filter (Millipore S2GPU05RE) to remove any cells, and the filtrate saved at 4 ºC (fraction A, phage prep). 15 mL of fraction A was filtered again using a 50 kDa filter (Millipore UFC905024) to remove any phages, and the ∼14.8 mL filtrate saved (fraction B, secretome without phages). ∼7.8 mL of fraction B was filtered again using a 3 kDa filter (Millipore UFC900324) to separate proteins larger and smaller than 3 kDa, and both the concentrate (fraction C, >3 kDa, ∼600 µL each) and the filtrate (fraction D, <3 kDa, ∼7 mL each) were saved (see Fig. 4c). The volumes of the fractions from each culture, listed in Table S3, were used to estimate the viscosity of the fractions (see Note S7). All four conditioned media fractions from the three strains were used as background to do a phage infection CFU assay using the AmpΦ phage. Briefly, 84 µL of the conditioned media fraction was mixed with 18.7 µL of the AmpΦ phage (undiluted concentration 8.8×10^3^ CFU mL^-1^) at RT, followed by adding 84 µL of ER2738 receivers (previously grown to a spectrophotometer OD_600_ of 2.2 and cooled down) and incubation at RT for 20 min. Next, 50 µL (26.7%) of the above infection reactions were diluted in 150 µL fresh LB and spread on LBA plates (200 µg mL^-1^ ampicillin) that were incubated at 37 ºC overnight and the colonies counted the next day (Fig. 4d).

### Effect of intracellular gIII in receivers on phage infections

Three colonies each of four receiver strains (ER2738, ER2738_HΔgIII, ER2738_H, ER2738_gIII-KanΦ) were grown to spectrophotometer OD_600_ of 1-1.3, cooled down and used for CFU assays using the AmpΦ phage (pSEVA19_M13ps, 1.85×10^4^ CFU mL^-1^), as described previously (90 µL cells + 10 µL phage prep). Colonies were counted the next day (Table S3) and plotted (Fig. S24).

### Secretome analysis

Protein content in the conditioned media fractions C from above (strains TOP10_H_KanΦ, TOP10_Gx_Kx, and TOP10_Gx_gIII-KanΦ) was quantified using Bradford assay (26, 26, and 32 ng µL^-1^, respectively), concentrated from 250 µL to ∼50 µL using a SpeedVac (Thermo Savant DNA120), and sent for shotgun proteomics analysis to the PAPPSO platform (http://pappso.inrae.fr/en/). At PAPPSO, 7.5 µL (∼1 µg) each of the samples was short run on an SDS-PAGE and excised. In-gel trypsin digestion was performed on the excised gel-bands from the SDS-PAGE, and the peptides extracted and analysed by LC-MS/MS according to the platform’s protocol^78^, adapted for the NanoElute system. The MS/MS spectra obtained were searched against NCBI databases (*Enterobacteria phage* NC_003287.2 and *Escherichia coli* NC_000913.3) using X!TandemPipeline (version 0.4.18)^79^, the open search engine developed by PAPPSO (http://pappso.inrae.fr/en/bioinfo/xtandempipeline/). Precursor mass tolerance was 20 ppm and fragment mass tolerance was 0.02. Data filtering was achieved according to a peptide E-value < 0.01, protein E-value < 10e-4, and to a minimum of two identified peptides per protein. Sample from TOP10_Gx_gIII-KanΦ had more proteins (494) detected than from TOP10_H_KanΦ (214) or TOP10_Gx_Kx (196). The normalised abundance (expressed as spectral counts) of each protein was determined per sample. The relative abundance for M13 pIII is plotted in Fig. 4g.

### Statistical Tests

We determined whether different conditioned media led to significantly different experimental outcomes (Fig. 4d). To compare two experimental outcomes (CFU counts) with each other, we used Welch’s t-test. To compare experimental outcomes to model predictions, we used the one-sample t-test. p-value of ≤0.005 is considered significant. Statistical tests were performed in Python (version 3.9.10) using the scipy.stats library (scipy version 1.5.4).

### Modelling

We summarise here the main modelling decisions. For a detailed discussion of the model, we refer the reader to **Note S8** in the supplementary material.

Conversion between OD_600_ measurements of the 96-well plate reader and the spectrophotometer was performed by linear regression (Python 3.9.10, sklearn toolkit version 0.24.1, see **Note S2** and **Fig. S4**). Conversion of OD_600_ measurements to cell densities for receiver and sender strains was by cubic fits (Python 3.9.10, scipy.optimize.curve_fit version 1.5.4, see **Note S2** and **Fig. S5**) of CFU counts over OD_600_ measurements.

We used Chemical Reaction Networks (CRNs) that describe the dynamics of a system by species and reactions that act upon them (**Box S3**). The CRN models were simulated with a custom simulation framework (https://github.com/ROBACON/BCRNsim) in Python (version 3.9.10) that generates SBML models and internally runs CopasiSE (version 4.34, build 251) to obtain stochastic (by the direct method) and ODE (by the LSODA solver) simulations. Simulations were parallelised using joblib, and the simulation data was post-processed and plotted in Python. Parameterisation of the initial CRN model and subsequent extensions began with a basic growth model from corresponding experimental data, which was then iterated on by adding new parameters and reactions to the model (phage secretion, infection, antibiotics, inhibition via external pIII) to fit more involved experimental setups. Fitting was done by particle swarm optimisation in Python and, if needed, adjusted manually. We next summarise the iterative model build-up (for details see **Note S8**).

We resorted to a linear resource consumer model with two resources to model the experimentally observed two-phase exponential growth of senders and receivers (**Note S3, Box S3**, and **Fig. S8**). Starting from the growth models, we first integrated phage secretion into the sender model (**Fig. 1**). We then detailed the receiver model by adding infection kinetics **(Fig. 1f-h**). The infection model uses a reaction with first-order kinetics for infection of receivers by phages (**Fig. 1g&i**) and two infectability states with different infection rates for a receiver (**Fig. 1h&j**). We further expanded the receiver CRN model by uptake and decay reactions induced by the antibiotics (e.g., kan). The newly introduced parameters for antibiotic uptake and death were calibrated with plate reader runs and CFU counts (**Fig. 1**, see **Methods** for experimental setup). An analytic first-order kinetics infection model was derived for the case where bacterial growth can be neglected and used to relate CFU and PFU counts (**Note S6**). CRN sender and receiver models were then combined into a single CRN to model end-to-end communication of senders to receivers, including phage secretion, infection, and immunity to different antibiotics (tet, gent, kan). The newly introduced parameters of tet and gent uptake and death were calibrated via plate reader runs (**Fig. 3**, see **Methods** for experimental setup). To model and confirm inhibition via external pIII in conditioned media, we resorted to a simplified, static first-order kinetics infection model (**Fig. 4d**). In a second step, the full receiver CRN model from **Fig. 3** was extended by inhibition via external pIII. Inhibition was assumed to follow first-order kinetics and block entry of phages. The newly introduced parameters were fitted with plate reader experiments (**Fig. 4e**, see **Methods** for experimental setup). The simplified wild-type M13 model in the gut (**Fig. 5**) was obtained by combining secretion and infection kinetics of senders and receivers (so far associated with separate cell types), allowing the same cell to have both functions. Since newly introduced additional parameters like the pIII secretion rate and the reduced out-flow rates of cells due to adhesion to the gut mucus could not be fitted with experimental data, we varied these parameters over large ranges to test for the robustness of our predictions.

## Acknowledgements

We thank Jean-Loup Faulon for his support and generous access to various laboratory equipment, and Marie-Agnès Petit for access to the UVP ColonyDoc-It. We thank Alfonso Jaramillo for the kind gift of bacterial strains (TOP10, TG1, SAJ635) and plasmids (PAJ297, PAJ156), and Vijai Singh for help in constructing PAJ156. We thank Hadi Jbara, Roman Luchko, Abhinav Pujar, and Tom Zaplana for technical assistance. We thank Sayantan Bose, Alfonso Jaramillo, and Marie-Agnès Petit for valuable comments and discussions. We thank Lydie Oliveira Correia and Aaron Millan-Oropeza (PAPPSO) for the proteomics analyses, which were performed on the PAPPSO platform (http://pappso.inrae.fr/en/) supported by INRAE (https://www.inrae.fr/en), the Ile-de-France regional council (https://www.iledefrance.fr/education-recherche), IBiSA (https://www.ibisa.net), and CNRS (http://www.cnrs.fr). We thank the Max Planck Institute for Informatics (MPI-INF) for access to cluster computing time. We acknowledge support from the Digicosme working group HicDiesMeus, Ile-de-France (IdF) region’s DIM-RFSI (project COMBACT), INS2I CNRS (project BACON), Université Paris-Saclay’s STIC department (project DEPEC MODE), and INRAE’s MICA department (project PHEMO). This research was funded in part by the National Research Agency (ANR) under the DREAMY project (ANR-21-CE48-0003).

## Author contributions

M.F., T.N., and M.K. conceived the study.

A.Pat., and M.K. designed the wet-lab experiments.

A.Pat., C.H., A.Pan., and M.K. performed the wet-lab experiments. C.H., M.F., and T.N. built and simulated the computational models. A.Pat., M.F., T.N., and M.K. analysed the data.

M.F., T.N., and M.K. acquired the funding.

M.F., T.N., and M.K. wrote the manuscript with contributions from all authors. All authors read and approved the final manuscript.

